# Bacterial but not protist gut microbiota align with ecological specialization in a set of lower termite species

**DOI:** 10.1101/083683

**Authors:** Lena Waidele, Judith Korb, Sven Küenzel, Franck Dedeine, Fabian Staubach

## Abstract

The role of microbes in adaptation of higher organisms to the environment is becoming increasingly evident, but remains poorly understood. Protist and bacterial microbes facilitate that lower termites thrive on wood and are directly involved in substrate break down. During the course of evolution lower termites adapted to different diets and lifestyles. In order to test whether there are changes of the termite gut microbiota that co-occur and hence could be related to diet and lifestyle adaptation, we assessed the bacterial and protist communities in a multispecies framework profiling three wood-dwelling and two foraging lower termite species using 16S and 18S rRNA gene amplicon sequencing. Termites were kept under controlled conditions on the same diet to minimize environmental effects on their gut microbiota. We found that protist communities group according to host phylogeny while bacterial communities group according to lifestyle. The change from the ancestral wood-dwelling to a foraging lifestyle coincides with exposure to more diverse and higher concentrations of pathogens as well as a more diverse diet. Accordingly, we identified bacteria that are associated with foraging termites of the genus *Reticulitermes* and could function as probiotics or be metabolically important on a more diverse diet. Furthermore, protist and bacterial diversity are correlated, suggesting not only that many termite gut bacteria are associated with protists, but also suggesting a role of protist diversity in the evolution of bacterial diversity in the termite gut or vice versa.

## Introduction

The importance of microbes in the evolution of higher organisms is starting to be realized (Feldhaar, 2011; McFall-Ngai et al., 2013; Wang et al., 2015). Metazoan evolution is not only driven by pathogenic microbes. Instead, microbes often are important facilitators of adaptation of higher organisms to the environment (McFall-Ngai et al., 2012; Douglas, 2015). The evolution of organisms is often so strongly intertwined with that of their associated microbes that Zilber-Rosenberg and Rosenberg (2008) suggested the host organism and its associated microbes should be treated as an evolutionary entity, the holobiont. However, most studies that aim at shedding light on the mechanisms of adaptation still focus on the host organisms all but ignoring its associated microbial communities.

The gut microbial communities of wood-feeding roaches and termites facilitated their ability to thrive on a wood diet. Wood is difficult to digest and poor in nitrogen. The ability to thrive on this resource has intrigued scientists for decades (e.g. Cleveland, 1923, 1925, Breznak et al. 1973). This interest was further spurred by the perspective to leverage insights into the process of lignocellulose break down in termites for the generation of biofuel (Scharf, 2015). Today, we can rely on extensive insights into the processes of lignocellulose break down and nitrogen fixation in the termite gut (Breznak et al., 1973; Warnecke et al., 2007; Hongoh et al., 2008b, 2008a; Brune and Dietrich, 2015; Ohkuma et al., 2015). For the breakdown of lignocellulose, lower termites depend on protists in their gut (Cleveland, 1923, 1925). These protists belong to the order Oxymonadida (phylum Preaxostyla), which are specific to termite and wood-feeding roach guts, and the phylum Parabasalia. The protists are transmitted between colony members and from parent to offspring via proctodeal trophallaxis. This vertical transmission contributes to patterns of cospeciation between protist and termite host (Noda et al., 2007; Desai et al., 2010). In return, most of the protists live in symbioses with ecto- and endosymbiotic bacteria. The bacterial termite gut microbiome consists of these protist-associated symbionts as well as bacteria freely living in the gut (Bauer et al., 1999; Breznak, 2002). Although vertical transmission of bacteria between termites is not as strict as for the mostly anaerobic protists (Noda et al., 2009), cospeciation can occur (Noda et al., 2007; Ohkuma, 2008; Ikeda-Ohtsubo and Brune, 2009; Desai et al., 2010).

The microbial communities of termites share a common origin with the microbial communities of cockroaches (Ohkuma et al., 2009; Schauer et al., 2012; Dietrich et al., 2014). Since the evolutionary split of termites from cockroaches, lower termites have diversified and adapted to a variety of environments (Eggleton and Tayasu, 2001), lifestyles (Abe, 1987), and diets (Donovan et al., 2001). Given the direct involvement of termite gut microbes in food digestion, the microbiota could play a role in adaptation to new diets (Zhang and Leadbetter, 2012). Prokaryotes that are beneficial for the termite host on a new diet can be acquired from the environment. The presence of such newly acquired microbes may indeed be reflected by the many taxon specific gut bacteria found in termites (Hongoh et al., 2005; Dietrich et al., 2014; Rahman et al., 2015; Tai et al., 2015). As a consequence of the acquisition of new bacteria that are beneficial under new dietary conditions, the termite associated microbial communities would change according to host diet. In fact, it has been shown that both, diet and evolutionary history, shape the gut microbiota of termites (Boucias et al., 2013; Dietrich et al., 2014; Rahman et al., 2015; Tai et al., 2015), with a larger impact of diet in higher termites (He et al., 2013; Mikaelyan et al., 2015a; Rossmassler et al., 2015). While the phylogenetic imprint on microbial communities is evident, it is not completely clear from these studies what leads to the diet related differences in microbial communities. These differences could be driven by environmental microbes that are ingested and passage the gut. Such passengers do not necessarily form any evolutionary relationship with termites. However, such a relationship would be expected for microbes that play a role in diet adaptation. In order to effectively disentangle passaging microbes from microbes that persist in the termite gut, potentially forming a stable relationship, it is necessary to control the environment of the termite host including the diet.

Furthermore, termites that change their lifestyle from wood-dwelling (single piece nesting according to Abe 1987) to foraging outside the nest (separate and intermediate nesting type according to Abe 1987), are expected to encounter more diverse and a higher load of microbes. Most foraging termite species are exposed to microbe rich soil (4x10^6^ colony forming units, Vieira and Nahas, 2005), carrying on average 5000 times more microbes than even the nests of damp wood-dwelling termites (800 colony forming units, Rosengaus et al. 2003). As a consequence microbial diversity should be higher in foraging species. Moreover, an increase in bacterial community diversity should be more pronounced than an increase in protist diversity because bacteria are more readily acquired from the environment when compared to anaerobic protists. The foraging lifestyle also opens the termite colony to invasion by fungal and bacterial pathogens (Korb et al., 2012, 2015) These pathogens exert selection pressures on the hosts that have led to a variety of defensive strategies (Rosengaus et al., 2011). However, the possibility that microbes themselves play a role in the defense of social insect colonies as probiotics or defensive symbionts has largely been neglected. This is surprising because defensive symbionts are known from many other insects (e.g. Kaltenpoth et al., 2005).

In this study we sought to test whether an evolutionary switch in host lifestyle and diet co-occurs with a change in microbial communities of lower termites. Furthermore, we intended to identify microbes that correlate in presence or abundance with this evolutionary switch, and hence could be functionally related to evolutionary change. The third objective was to test whether the foraging termites in our study carry an increased microbial diversity, specifically bacterial diversity. Therefore, we analyzed the microbial communities of five termite species that were kept under common conditions. These species comprised two species of wood-dwelling kalotermitids (*Cryptotermes secundus*, *Cryptotermes domesticus*), a wood-dwelling rhinotermitid (*Prorhinotermes simplex*), and two species of rhinotermitids that switched from the ancestral wood-dwelling lifestyle to foraging (*Reticulitermes flavipes*, *Reticulitermes grassei,* Figure 1). Because there can be substantial variation of microbial communities between different colonies (Hongoh et al., 2005, 2006; Benjamino and Graf, 2016), we analyzed four to eight replicate colonies per species.

**Figure 1:**
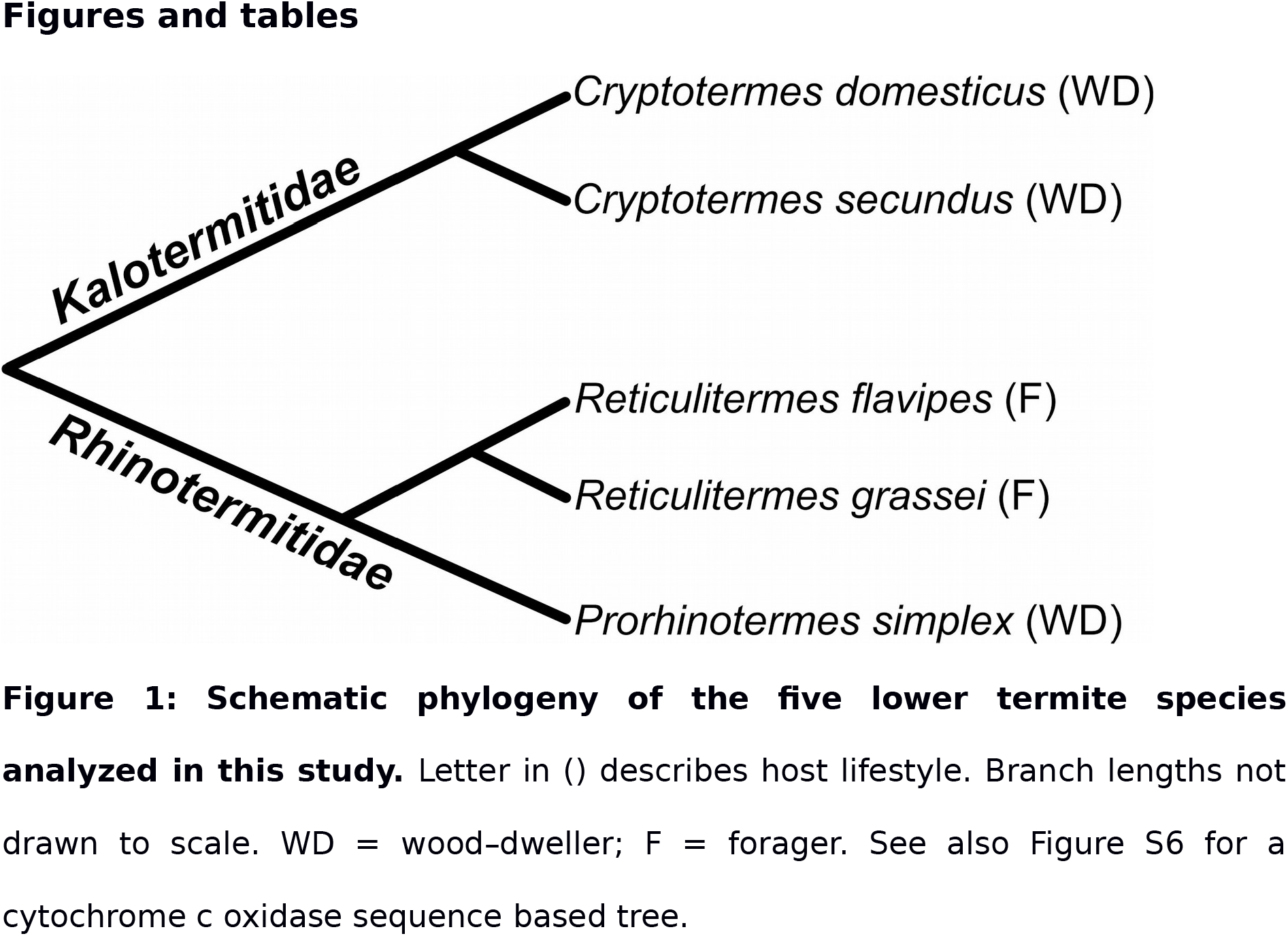
Schematic phylogeny of the five lower termite species analyzed in this study. Letter in () describes host lifestyle. Branch lengths not drawn to scale. WD = wood–dweller; F = forager. See also Figure S6 for a cytochrome c oxidase sequence based tree.

## Results

We used the Illumina MiSeq platform to sequence ~250 base pairs (bp) of the 16S rRNA gene to cover the v4 region for bacterial community profiling of the gut microbiota. Protist communities were profiled using a set of 18S rRNA gene specific primers of our own design that targeted the parabasalid protists(see Materials and Methods). Oxymonads were excluded from the analysis because they are difficult to target (e.g. Heiss and Keeling, 2006) and escaped reliable illumina sequencing compatible amplification (data not shown). All termites were kept on *Pinus radiata* wood from the same source. After quality-filtering, our bacterial and parabasalid datasets contained 1 676 707 and 2 747 988 gene sequences respectively (Table S1).

### Diversity of microbial communities

OTU (Operational Taxonomic Unit) based analysis of these sequences revealed that bacterial as well as parabasalid diversity was higher in the foraging species that we analyzed (Wilcoxon test on Shannon diversity: *P* = 0.005, and *P* = 0.03 respectively, Table 1). Rarefaction curves representing the microbial diversity in each termite species can be found in Supplementary Figure S1. Higher microbial diversity is expected in foraging species if we assume that species frequently leaving their nests to forage have a higher probability of encountering and acquiring new microorganisms.

**Table 1:** Average Shannon – diversity indices of bacterial and protist communities based on 97% sequence similarity OTUs. n = number of replicate colonies for each species; H = Average Shannon-Diversity; SD = Standard deviation; *P* = p-values of Wilcoxon test on Shannon-Diversity wood- dweller vs. forager.

|  | *C.*<br>*domesticu*<br>*s* | *C.*<br>*secundu*<br>*s* | *P.*<br>*simple*<br>*x* | *R.*<br>*flavipe*<br>*s* | *R.*<br>*grasse*<br>*i* | wood-<br>dweller | forager | *P* |
| --- | --- | --- | --- | --- | --- | --- | --- | --- |
| n | 5 | 8 | 7 | 5 | 4 | 20 | 9 |  |
| Bacterial community |  |  |  |  |  |  |  |  |
| OTUs | 450 | 509 | 490 | 697 | 559 | 481 | 636 | 0.005 |
| H | 3.56 | 3.42 | 3.14 | 4.3 | 3.72 | 3.37 | 4.04 |  |
| SD | 0.25 | 0.3 | 0.45 | 0.25 | 0.87 | 0.37 | 0.64 |  |
| Parabasalid community |  |  |  |  |  |  |  |  |
| OTUs | 31 | 21 | 21 | 36 | 36 | 23 | 36 | 0.03 |
| H | 1.15 | 1.37 | 0.7 | 1.63 | 1.23 | 1.08 | 1.45 |  |
| SD | 0.52 | 0.2 | 0.02 | 0.1 | 0.37 | 0.4 | 0.32 |  |

Furthermore, the diversity of parabasalid protists and bacteria was correlated across termite species (Pearson’s product – moment correlation coefficient: *P* = 0.003, r^2^ = 0.52, Figure 2). This correlation supports the notion that many bacteria present in the gut of lower termites are directly associated with protists. However, the bacterial diversity found in samples from the foraging *Reticulitermes* was higher than predicted by the regression from protist diversity of wood-dwelling species (ANOVA: *P =* 0.005, see Table S2 for linear models and tests performed). This suggests that a larger proportion of the bacterial diversity in *Reticulitermes* guts is either associated with oxymonads or does not live associated with protists when compared to wood-dwellers.

**Figure 2:**
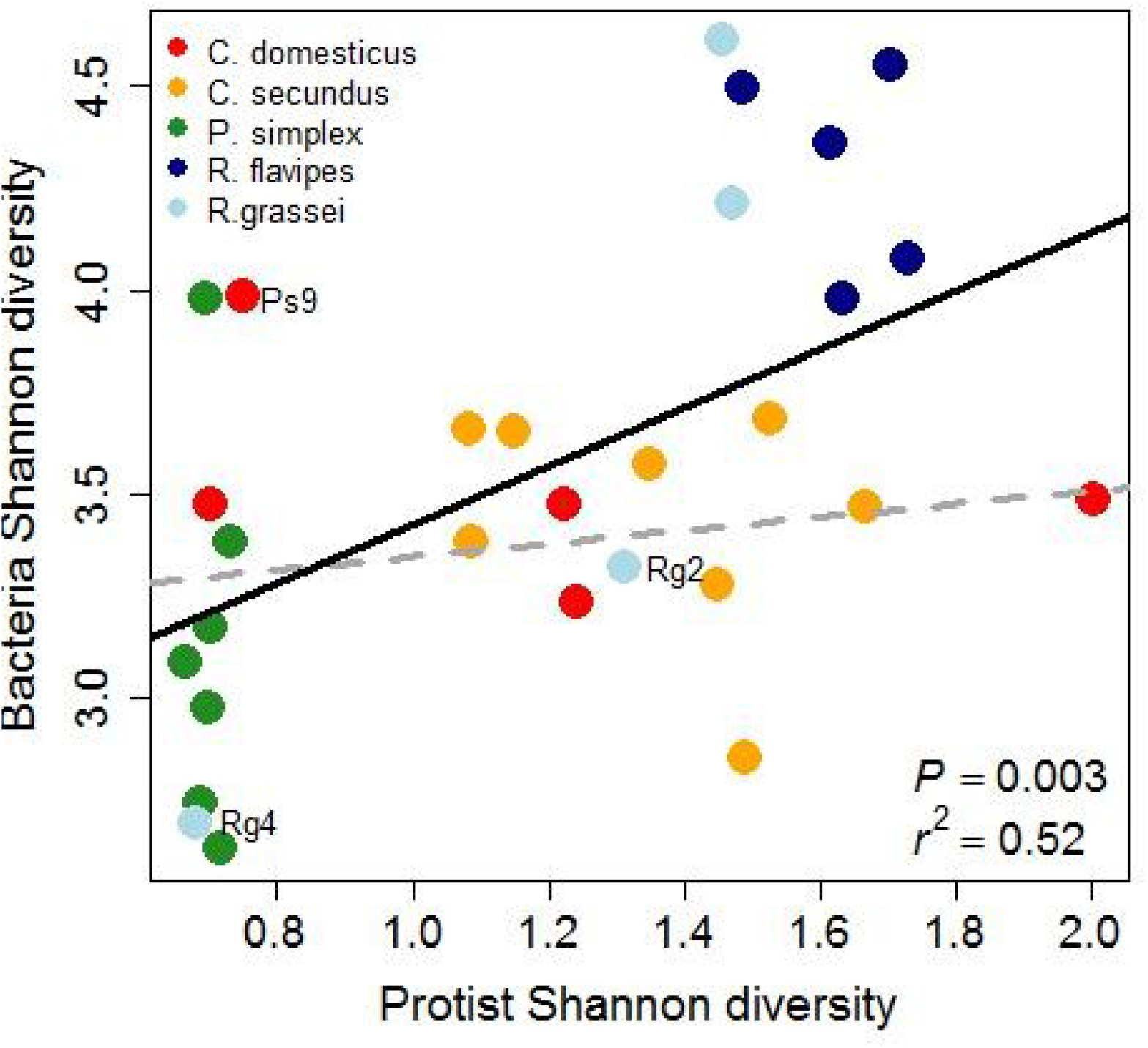
The Shannon diversities of protists and bacteria are correlated across species. Each dot represents a termite colony. The solid regression line is based on all samples, while the dashed line is based on wood- dwellers only. Colonies with unusual diversity patterns were marked (Ps9, Rg2, and Rg4, see discussion).

*Reticulitermes grassei* samples Rg2 and Rg4 as well as *P. simplex* sample Ps9 showed unusual bacterial diversity and community profiles (see discussion). We performed all diversity comparisons with and without these samples and found no qualitative difference in the results. Excluding these samples would increase average *R. grassei* bacterial community diversity, decrease *P. simplex* community diversity and lead to lower p-values in all tests shown in Table 1.

### Composition of microbial communities

The correlation of bacterial and protist community diversity that we found suggests that many of the bacteria are associated with protists. Indeed, taxonomic classification of 16S rRNA gene sequences revealed many protist associated bacterial taxa (Figure 3A, B). The four most common bacterial phyla in our samples were *Bacteroidetes* (40.6%), *Spirochaetes* (21.9%), *Proteobacteria* (12.4%), and *Elusimicrobia* (9.1%). Of these phyla the *Bacteroidetes*, *Spirochaetes*, and *Elusimicrobia* contain members known to be important ecto- or endosymbionts of lower termite associated protists (Noda et al., 2005, 2006; Ohkuma, 2008; Strassert et al. 2010).

**Figure 3:**
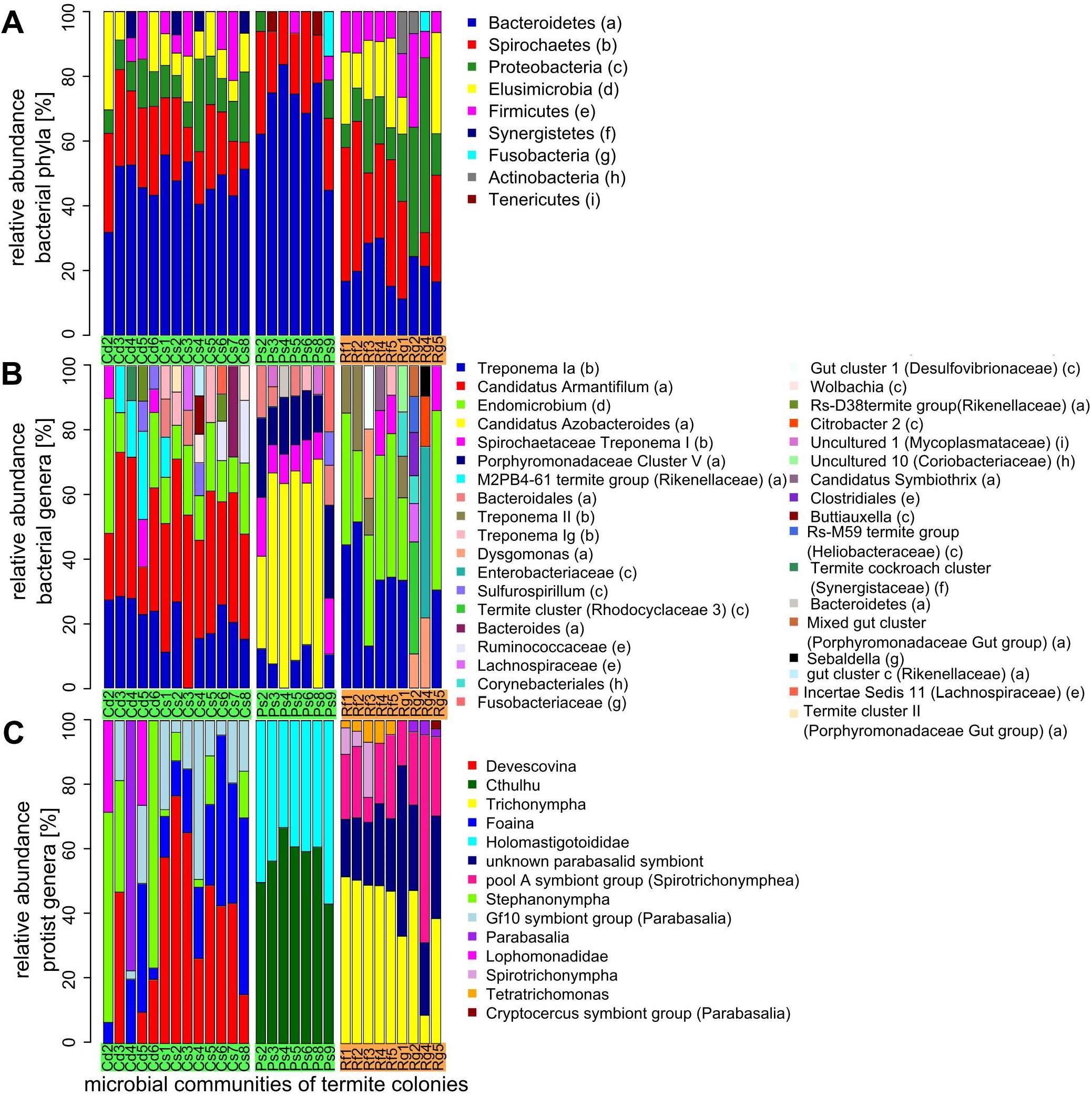
Relative abundance of bacterial and protist taxa as assessed by taxonomic profiling of 16S – and 18S rRNA gene sequences. Legends are sorted by abundance beginning with the most abundant taxa (A) Relative abundance of bacterial phyla. Phyla with a relative abundance <5% were removed for clarity. (B) Relative abundance of bacterial genera. Sequences that could not be classified to genus level with sufficient bootstrap support were labeled with the next higher taxonomic rank that could be assigned with sufficient support. Genus level classifications with unconventional names (e.g. “Gut cluster 1”) are followed by family classification in parentheses. Letters in parentheses correspond to phyla in A. Genera of abundance <5% have been removed for clarity. (C) Relative abundance of protist genera. See B for non genus level classifications. Genera with an abundance <2% have been removed. Cd = colonies of *C. domesticus*, Cs = *C. secundus*, Ps = *P. simplex*, Rf = *R. flvipes*, Rg = *R. grassei*.

The phylum *Bacteroidetes*, contains the genus *Candidatus* Armantifilum, which was common in all *Cryptotermes* colonies (14.7 – 53.7%). *Candidatus* Armantifilum devescovinae is a filamentous bacterium that can be found attached to the surface of *Devescovina* (Lophomonadidae) flagellates and cospeciates with them in kalotermitids (Desai et al., 2010). Close relatives of *Candidatus* Armantifilum are associated with *Snyderella tabogae* (Lophomonadidae) in *Cryptotermes longicollis (*Desai and Brune, 2012). Accordingly, we found lophomonadids in all *Cryptotermes* colonies (Figure 3C).

*Elusimicrobia* are well known endosymbionts of many termite-gut flagellates (Ikeda-Ohtsubo et al., 2007; Brune, 2012). Sequences from a prominent member of the *Elusimicrobia, Endomicrobium* were detected in all samples (0.3 – 27.8%). *Endomicrobia* are endosymbionts of *Trichonympha* (Ikeda-Ohtsubo and Brune, 2009) which we found in *Reticulitermes* (8.7 – 51.3%). Other *Endomicrobia* are symbionts of the *Cristamonadida* (Desai et al., 2010), of which we found *Devescovina* (0.2 – 76.2%), *Stephanonympha* (0 – 75.0%) and *Foaina*- species (0.6 – 54.2%) in *Cryptotermes*. Interestingly, a small proportion of sequences from the *Endomicrobiaceae* were found in the colonies of *P. simplex* (0.4 – 3.0%). A protist host for *Endomicrobia* has not yet been described in *P. simplex* and Tai *et al.* (2015) found no evidence for *Endomicrobia* in *P. simplex.* However, our sampling of 16S rRNA gene sequences was 8-10 times deeper than that of Tai et al. (2015) providing more power to detect rare microbes. 2 724 of 3 459 (86%) of the sequences from *Endomicrobia* found in *P. simplex* clustered into three OTUs (OTU 90, OTU 252, and OTU 316) that were specific to *P. simplex.* Hence, they are unlikely to represent contamination from non-*P. simplex* samples. These three OTUs are closely related and cluster phylogenetically with sequences from defaunated *R. santonensis* as well as a sequence from a *Zootermopsis nevadensis* flagellate suspension (Figure S2).

*Spirochaetes,* which are characteristic of termite guts, where they can be free living or live as ectosymbionts of protists (Breznak, 2002; Wenzel et al., 2003; Noda et al., 2003), were found in all samples (9.1 – 50.5%) except the abnormal sample Rg2.

Sequences from the spirochaete *Treponema Ia* fell into several distinct OTUs that were termite taxon specific (e.g. OTUs 2, 3, and 33, see S5). Many other bacterial genera contained termite taxon specific OTUs. For example, sequences from *Endomicrobium* fell into several taxon specific OTUs (7, 12, and 20). *Candidatus* Armantifilum (OTUs 1, 31, and 48) separated into termite genus specific OTUs as well. The prevalence of host specific OTUs suggests that the microbial communities of the termite species analyzed here are distinct. At the same time, the relatedness of these OTUs from different termite taxa, as reflected by their common classification into a single genus, supports the notion that the members of these communities often share a phylogenetic history (Schauer et al., 2012; Dietrich et al., 2014).

### Analysis of microbial community differences and similarities

The phylogenetic relationship between bacteria from different termite species can be incorporated into community distances by analyzing environment specific phylogenetic branch lengths of all community members (Lozupone and Knight, 2005). We were especially interested in the question whether similarities in microbial community composition reflect host diet and lifestyle. Assuming that microbes that are ecologically relevant for the host should not be rare, we used the weighted Unifrac metric (Lozupone et al., 2007) for our analysis.

The bacterial communities of the wood-dwelling species *C. domesticus, C. secundus* and *P. simplex* clustered separately from the bacterial communities of the foraging *Reticulitermes* (Figure 4A) and together with the bacterial communities of the wood-dwelling cockroach *Cryptocercus* (Figure S5), which is a sister taxon of termites and also wood-dwelling. In contrast, the parabasalid protist communities clustered strictly according to termite host family (Figure 4B). Interestingly, the parabasalid communities of *R. flavipes* and *R. grassei* were distinct (Approximately Unbiased support = 93%), while there was no consistent difference between the protist communities of *C. secundus* and *C. domesticus.* This might in part be due to the high variability of protist communities within *C. domesticus.* The relative sequence abundance of the five most common protists in *C. domesticus* varied substantially (*Devescovina*: 0.4% – 77.1%, *Stephanonympha*: 0.3% - 75.0%, *Foaina*: 0.6% - 39.0%, unclassified lophomonadid: 0.07% - 28.3%, unclassified parabasalid: 1.4% - 23.3%). Samples Cd2 and Cd6 had similar protist communities that were somewhat distinct from the other *Cryptotermes*. These samples shared a very low relative abundance (1.7% and 1.4%) of an unclassified genus of the Gf10 symbiont group that is common in the other *Cryptotermes* samples (red in Figure 3C). They also share a high prevalence of *Stephanonympha* (62.8% and 75.0%).

**Figure 4:**
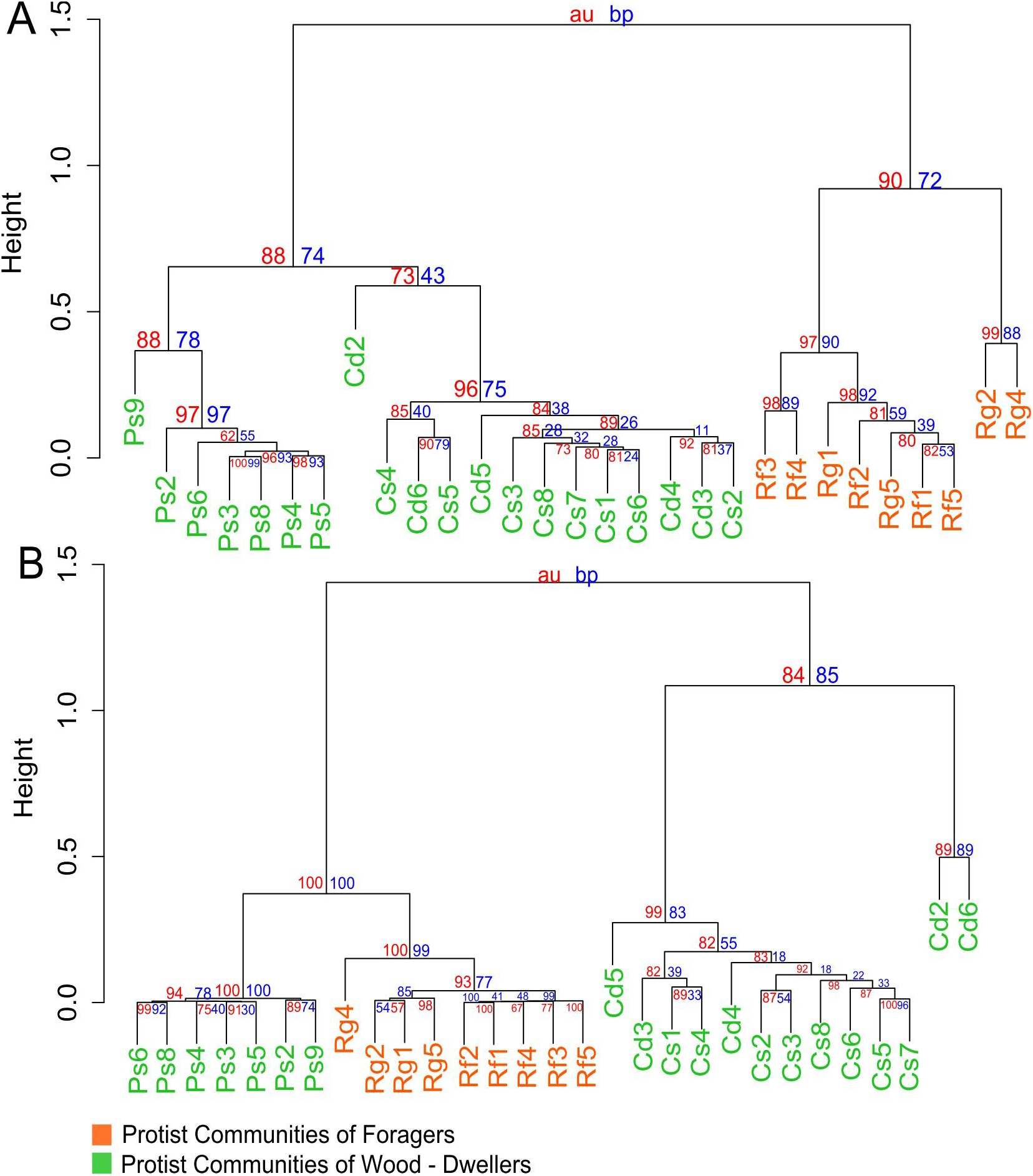
Cluster dendrograms based on weighted UniFrac community distances. (A) Cluster dendrogram based on bacterial community distances. (B) Cluster dendrogram based on protist community distance. Cd = colonies of *C. domesticus*, Cs = *C. secundus*, Ps = *P. simplex*, Rf = *R. flavipes*, Rg = *R. grassei*. blue numbers = bootstrap probability; red numbers = approximate unbiased probability.

### Bacteria specifically associated with foraging *Reticulitermes*

The clustering of foraging *Reticulitermes* bacterial communities suggests that there are bacteria specifically associated with *Reticulitermes.* In order to detect these bacteria, we used an Indicator Species Analysis (Dufrêne and Legendre, 1997). Foraging *Reticulitermes* and wood-dwelling termites were treated as two different habitats and bacterial 97% identity OTUs as species. A full list of Indicator OTUs is available in Table S10.

*Reticulitermes* indicator OTUs that are more closely related to environmental bacteria or bacteria found in non-dictyopteran hosts than to OTUs found in *P. simplex* could be recently acquired by *Reticulitermes*. In order to shed light on which bacteria are closely related to *Reticulitermes* indicator OTUs, we constructed a phylogenetic tree. This tree included representative sequences from the indicator OTUs, all OTUs for which we found at least 100 sequences in *P. simplex* or *Cryptotermes,* and all sequences in DictDB (Mikaelyan et al., 2015b) (Figure S4). The Newick formatted phylogeny can be found in S8.

Table 2 contains a list of *Reticulitermes* indicator OTUs that do not have *P. simplex* OTUs nor bacteria from other wood-dwellers amongst their closest relatives and contain more than 1 000 sequences. Close relatives of all of these indicator OTUs were found in association with *Reticulitermes* in previous studies, corroborating our results. Close relatives of several indicator OTUs came from the environment, non-dictyopteran hosts or other foraging termites.

**Table 2:**
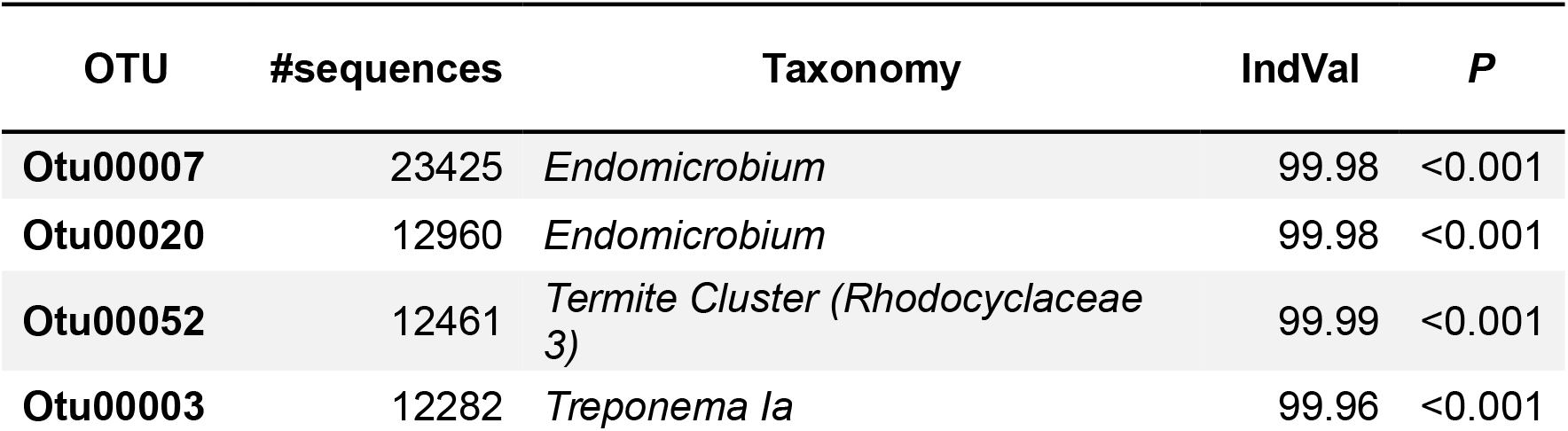
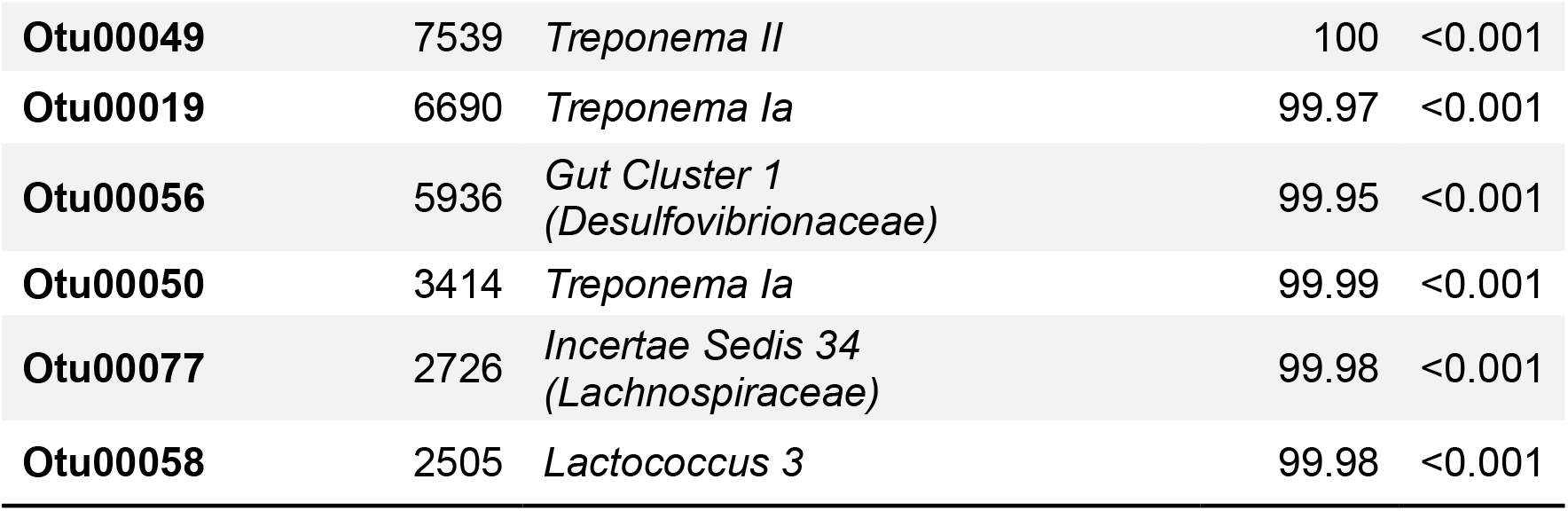
Bacterial Indicator OTUs of *Reticulitermes*. Shown are indicator OTUs with an indicator value larger than 99.9 and *P* < 0.001 that comprise at least 1,000 sequences and have no known close relatives in wood-dwellers. IndVal = Indicator value; DictDb ID refers to the label set in DictDb_v3.

## Discussion

Profiling bacterial and parabasalid protist communities, we found (i) that protist and bacterial diversity correlate, (ii) evidence for changes in community composition and diversity that correlate with an evolutionary switch from wood-dwelling to foraging, (iii) candidate microbes that are specific to foraging *Reticulitermes*. Furthermore, we found several samples that showed unusual community profiles that require discussion.

### Unusual community profiles and intraspecific variation in microbial communities

Three samples showed unusual community profiles (Ps9, Rg2, and Rg4). More than half of the sequences from Rg4 are comprised by OTU 16. OTU 16 was classified as a member of the Enterobacteriaceae. Many enterobacteria are insect pathogens (Grimont and Grimont, 2006; Galac and Lazzaro, 2011; Jurat-Fuentes and Jackson, 2012). Potential insect pathogens can often be found at low titers in natural host populations where they probably live as commensals (Staubach et al., 2013). This is also true for OTU 16 that comprises no more than 8% of the sequences in the other samples. However, these bacteria can reach high titers when entering the hemolymph (Galac and Lazzaro, 2011) causing a systemic infection. This renders it plausible that Rg4 contained a termite that was systemically infected with OTU 16.

The bacterial communities of Rg2 and Ps9 contained a smaller fraction of sequences from presumably protist associated bacteria (*Endomicrobium* and *Candidatus* Azobacteroides) than their conspecifics (1.2% instead of 10.1 - 27.4%, and 0.01% instead of 19.2% – 49.3%, respectively). This suggests that these samples might have had a lower protist titer. A low protist titer in termites is commonly caused by loss of protists from the hindgut during molts (Honigberg, 1970). We selected individuals that had filled guts and carefully excluded individuals that were close to molting indicated by their whitish appearance. However, these samples might have contained termites that have not fully reestablished their protist communities after molting. Because we carefully selected the termites for our study, it appears unlikely that the overall high variability in the protist communities of *C. domesticus* is purely a result of undetected molts. The variance in community composition between conspecific termite colonies that we and others (Hongoh et al., 2005, 2006; Benjamino and Graf, 2016) observed suggests that the use of replicate colonies is important.

### Bacterial and protist diversity

We presented, to our knowledge, the first report of a correlation between bacterial and protist community alpha diversity. Studies that analyze both, protist and bacterial communities in parallel, across termite species are still rare. In the two other studies, we are aware of, neither environmental nor dietary conditions were controlled (Tai et al., 2015; Rahman et al., 2015). As a consequence, the diversity patterns of termite associated microbial communities might have been affected by microbes merely passing through the gut. These passengers could have obscured the correlation of protist and bacterial diversity that we observed.

A correlation between protist and bacterial diversity does not only suggest that many bacteria are protist associated in the termite gut (Brugerolle and Radek, 2006). This correlation also implies a potential mechanism, how the high bacterial diversity in the termite gut (Colman et al., 2012; Jones et al., 2013) has evolved. A higher protist diversity offers bacteria a larger number of independent bacterial habitats that can be colonized. Because there is good evidence that the gut protists vertically transmit their bacterial symbionts (Noda et al., 2007; Ikeda-Ohtsubo and Brune, 2009; Desai et al., 2010) and can carry a single isolated strain of their symbionts (Ikeda-Ohtsubo et al., 2007; Zheng et al., 2015), these bacteria experience reduced gene flow that allows for the accumulation of genetic differences. Furthermore, the different environmental conditions, each protist species offers, exert selection pressures on their associated bacteria further promoting divergence via natural selection. In fact, the protists could serve as 'speciation reactors' for bacterial symbionts and facilitate the evolution of high bacterial diversity in termite guts. On the other hand, coadaptation between protists and bacteria could promote protist speciation (Dolan, 2001).

The bacterial diversity in the gut of *Reticulitermes* was higher than expected given parabasalid protist diversity of the other species. This implies that *Reticulitermes* carries either a higher proportion of oxymonad protist associated bacteria or more bacteria that live independently of protists. Because the proportion of oxymonad to parabasalid diversity is identical in *R. flavipes* (synonym *R. santonensis*) and *C. domesticus* (1:2, according to Yamin, (1979), non-protist associated bacteria are likely to contribute to the additional bacterial diversity in *Reticulitermes.*

Our bacterial diversity estimates for Reticulitermes are in concordance with estimates from Benjamino and Graf (2016) who sequenced the same fragment of the bacterial 16S rRNA gene. Their estimates of Shannon diversity range from 3.54 to 4.48 for different colonies and fit our estimate of 4.3+-0.25 well. A previous study that reached a similar sequencing depth of an overlapping region of the 16S rRNA gene also estimated similar Shannon diversities for *R*. *flavipes* of 3.92 (Dietrich et al., 2014). This is within one or two standard deviations of the diversity we calculated for *R. flavipes* and *R. grassei* (4.3+-0.25 and 3.72+-0.87, respectively). Rahman et al. (2015) estimated Shannon diversity of *Reticulitermes* to be somewhat higher (4.91-5.54). However, these authors included protists in their diversity estimates. In contrast, the bacterial diversity estimate in Tai et al. (2015) was lower with Shannon diversity for *R. flavipes* at 3.66 versus 3.92 in our study, as well as 0.81 versus 3.14 for *P. simplex*. This difference could result from our ~10X increased sampling depth. Shannon diversity is known to correlate with sampling depth (Preheim et al., 2013). Furthermore, Tai et al. (2015) used a masking approach to remove variable regions from their alignment. This masking can significantly reduce diversity and resolution (Schloss, 2010). Because we used SILVA (Pruesse et al., 2007) based DictDB (Mikaelyan et al., 2015b) for our alignments instead of Greengenes (DeSantis et al., 2006) as in Tai et al. (2015), this masking is not necessary (Schloss, 2010). The same arguments can explain the lower parabasalid protist diversity in Tai et al. (2015) (0.5 for *R. flavipes* compared to 1.63 in our study and 0.29 for *P. simplex* compared to 0.7 in our study). That our protist diversity estimates were higher than those obtained by visual inspection (Yamin, 1979) is not surprising since cryptic variation has been repeatedly documented from termite protists (Stingl and Brune 2003; Brugerolle and Bordereau, 2006; Zheng et al., 2015). Especially smaller protists often escape visual methods (Radek et al., 2014).

### Bacterial communities cluster according to lifestyle while protist communities cluster according to host phylogeny

As a consequence of the switch from the ancestral wood-dwelling lifestyle to foraging, *Reticulitermes* are not only exposed to more diverse microbial communities from outside the nest, but also experienced a change to a more diverse diet. *Reticulitermes* probably originated in inland habitats (Dedeine et al., 2016) where they feed on a variety of substrates (Waller, 1991). Their natural substrates include decaying wood (Amburgey, 1979), horse dung (Rahman et al., 2015), and they are even able to acquire nutrients from soil (Janzow and Judd, 2015). In contrast, *C. domesticus, C. secundus* and *P. simplex* are largely restricted to islands and coastal regions where they dwell on dead wood (Eggleton, 2000). These changes in lifestyle and diet align with a change of bacterial communities, but not protist communities. Therefore, microbes that are related to this change in lifestyle are more likely to be found amongst the bacteria. This result adds to the notion that bacterial communities are more variable over evolutionary time than protist communities (Noda et al., 2009; Tai et al., 2015).

A higher propensity of random gain and loss of bacteria in *Reticulitermes* that is fueled by exposure of foragers to more diverse microbes could leave the bacterial community of *Reticulitermes* distinct from that of wood-dwellers. However, *Prorhinotermes* have probably reverted back from foraging to wood-dwelling (Legendre et al. 2008, Bourgignon et al. 2015) leaving its ancestors exposed to similarly diverse microbes, and *Prorhinotermes* with a similar propensity for random change of microbial communities. Given similar potential for random change, invoking lifestyle related natural selection either driving the divergence of the *Reticulitermes*microbiota, or the convergence of the *Prorhinotermes* microbiota back to that of wood-dwelling species, or both, appears plausible to explain the distinct bacterial communities of *Reticulitermes*.

### Candidate bacteria that correlate with the change in lifestyle

DictDB (Mikaelyan et al., 2015b) allowed us to classify the majority (75.5%) of our bacterial sequences to the genus level and identify bacteria that correlate with the evolutionary switch in lifestyle from wood-dwelling to foraging in *Reticulitermes*. These bacteria come from diverse families. OTU 120 is a spirochaete from the genus *Treponema I*. Spirochaetes including *Treponema I* strongly correlate with diet in higher termites (Mikaelyan et al., 2015a; Rossmassler et al., 2015). Indicator OTU 307 was classified as *Stenoxybacter. Stenoxybacter* colonizes the hind gut wall of *R. flavipes* and contributes to acetate metabolism as well as oxygen consumption (Wertz and Breznak, 2007). Another interesting candidate is OTU 58 (*Lactococcus*). The genus *Lactococcus* has, amongst termites, so far only been reported from *Reticulitermes* and higher termites (Bauer et al., 1999; König et al., 2006; Mathew et al., 2012; Boucias et al., 2013; Yang et al., 2015). Lactococci are powerful probiotics in mammals (Ballal et al., 2015), fish (Heo et al., 2013) and arthropods (Maeda et al., 2013). The acquisition of probiotics would be an obvious evolutionary response to the increased pathogen exposure that is connected to the foraging lifestyle. Furthermore, *Lactococcus* might be metabolically important in *R. flavipes* as it directly or indirectly contributes to acetate production in termite guts (Bauer et al., 1999).

## Conclusion

We provide evidence that, even after removal of environmental noise through keeping different lower termite species under common conditions, an ecological imprint on their associate bacterial communities remains. Studies that control the termite environment include more species and span several evolutionary switches in lifestyle are necessary to generalize this pattern.

## Acknowledgements

We thank Jan Sobotnik for kindly providing *P. simplex* colonies and Andreas Brune for access to a pre-publication version of DictDB that made our life much easier. We thank Chris Voolstra and Volker Nehring for helpful comments on the manuscript. This work was funded by DFG grants STA1154/2-1 and KO1895/6-4.

## Materials and Methods

### Termite samples

All termites used in the experiments were kept under constant conditions (27°C, 70% humidity) and acclimated to the same food substrate (*Pinus radiata* wood) in our lab for at least six weeks prior to the experiment. This period is considered sufficient for termites and their gut microbes to adjust to a new diet (Huang et al., 2013) and lies far beyond the 24h passage time of food through the termite gut (Breznak, 1984). *C. domesticus* and *C. secundus* were collected in Darwin (in 2003, 2007 and 2011), Australia. *P. simplex* was collected in Soroa, Cuba (in 1964), and colonies of *R. flavipes* as well as *R. grassei* were collected on the Île d’ Oléron, France (in 2014). Morphological species identification was confirmed by sequencing of the mitochondrial cytochrome c oxidase subunit II (Figure S6 and supplementary methods) for most of the samples. The foraging species *R. flavipes* and *R. grassei* were additionally provided with sand to build their nests. In order to exclude a contribution of microbes or microbial DNA in the sand to the *Reticulitermes* associated microbiota, we applied the same DNA extraction, amplification, and sequencing procedure as for the termite guts to the sand. However, we failed to amplify the targeted fragments from the sand samples. This led us to conclude that the sand did not contribute to the bacterial nor protist community profiles.

In order to test whether the time since collection from the wild played a role in diversity and composition of the microbial communities we analyzed the microbial communities of *Cryptotermes* that have been in the lab for different periods of time (2004 – 2012). We could not detect any effect of the time under lab conditions on microbial community alpha diversity (bacteria: *P* = 0.99, r = 0.001; protists: *P* = 0.13, r^2^ = 0.21) nor composition and beta diversity (Supplementary Figure S3).

### DNA extraction, primer design and library preparation

DNA was extracted from the guts of three workers per colony using bead beating and chloroform precipitation (also see supplementary material and methods). We amplified the v4 region of the 16S rRNA gene with the bacteria specific primerset 515f (5’–GTGCCAGCMGCCGCGGTAA–3’) and 806r (5’– GGACTACHVGGGTWTCTAAT–3’) developed by Caporaso *et al.* (2012). Specific barcode combinations were added to the primer sequences following Kozich *et al.* (2013). In order to amplify a region of the 18S rRNA gene of parabasalid protists, we developed custom designed primers. Primer development was based on parabasalid sequences from SILVA SSU Ref Nr 123 that was extended by 18S rRNA gene sequences from protist taxa that we expected to occur in the termite species we analyzed according to Yamin (1979). These additional sequences were downloaded from ncbi and aligned using ClustalW (Larkin et al., 2007) in the BioEdit Sequence Alignment Editor version 7.2.5 (Hall, 1999). The resulting alignment was manually curated and searched for potential forward and reverse primers that would match as many protist sequences as possible but not termite 18S rRNA genes. The 18S rRNA gene alignment as well as the corresponding taxonomy reference - file are provided in the supplement (S7). The resulting primerset Par18SF: 5’-AAACTGCGAATAGCTC-3’; Par18SR: 5’- CTCTCAGGCGCCTTCTCCGGCAGT-3’, amplified ~250 bp. A detailed PCR protocol can be found in the supplementary material and methods section. Libraries were sequenced on an Illumina MiSeq reading 2x 250bp.

### Analysis

Data analysis was performed using MOTHUR version 1.33.3 (Schloss et al., 2009), following the MiSeq SOP. Our detailed MOTHUR script can be found in S9. For taxonomic classification of the bacterial 16S OTUs a representative sequence of each OTU was aligned to a termite specific database DictDb_v3 (Mikaelyan et al., 2015b), while 18S protist OTUs were aligned and classified using all parabasalid sequences from the SILVA SSU Ref Nr 123 database that was extended by our custom made alignment and taxonomy reference file (S7). In order to avoid coverage bias, bacterial and protist datasets were rarefied resulting in 32 824 and 22 285 sequences per sample, respectively. Statistical tests and data visualization were performed in R version 3.1.2 (2012). Phylogenetic trees necessary to apply UniFrac (Lozupone et al., 2007) as well as the trees including OTU representative sequences were generated with FastTree version 2.1.7 (Price et al., 2010). The resulting trees were visualized with Dendroscope 3.3.4 (Huson and Scornavacca, 2012). Trees based on Unifrac were generated using the UPGMA algorithm as implemented in the R PvClust package version 2.0.0 (Suzuki and Shimodaira, 2015) and bootstrapped 10 000 times with standard settings. Indicator species were obtained following the method by (Dufrêne and Legendre, 1997), as included in MOTHUR.

## Data availability

Sequence data is available at MGRAST (http://metagenomics.anl.gov/) under the IDs 4711469.3 – 4711607.3. Supplementary information is available at the journal website.

